# Phloem-Specific Translational Regulation of Soybean Nodulation: Insights from a Phloem-Targeted TRAP-Seq Approach

**DOI:** 10.1101/2025.04.09.647652

**Authors:** Jae hyo Song, Shin-ichiro Agake, Sayuri Tanabata, Yaya Cui, Li Su, Bruna Montes-Luz, Dong Xu, Gary Stacey

## Abstract

Soybean (*Glycine max*) root nodulation is a symbiotic process that requires complex molecular and cellular coordination. The phloem plays a crucial role not only in nutrient transport but also in long-distance signaling that regulates nodulation. However, the molecular mechanisms underlying phloem-specific regulation during nodulation remain poorly characterized. Here, we developed a phloem-specific Translating Ribosome Affinity Purification sequencing (TRAP-seq) system to investigate the translational dynamics of phloem-associated genes during nodulation. Using a phloem-specific promoter (*Glyma.01G040700*) combined with the GAL4-UAS amplification system, we successfully captured the translatome of soybean root phloem at early (72 hours post-inoculation, hpi) and late (21 days post-inoculation, dpi) nodulation stages. Differential expression analysis revealed dynamic translational reprogramming, with 2,636 differentially expressed genes (DEGs) at 72 hpi and 8,422 DEGs at 21 dpi. Gene ontology and pathway enrichment analyses showed stage-specific regulatory shifts, including early activation of ethylene and defense pathways and late-stage enhancement of nutrient transport and vascular development. Transcription factor analysis identified *GmbHLH121* as a key phloem-specific regulator of nodulation. Functional validation using RNAi knockdown and overexpression experiments demonstrated that *GmbHLH121* negatively regulates nodule formation, likely acting downstream of or independently from early nodulation signaling pathways. Additionally, we uncovered dynamic regulation of cell wall-modifying enzymes (*PME* and *PMEI*) in the phloem, implicating their role in modulating plasmodesmata permeability and facilitating symplastic connectivity during nodulation. Our findings highlight the critical role of phloem-mediated translational regulation in coordinating root nodulation, emphasizing the phloem as an active regulatory hub for long-distance signaling and symbiotic efficiency.

## Introduction

Soybean (*Glycine max*) phloem plays a pivotal role in facilitating symbiotic interactions essential for root nodulation and nitrogen fixation. Beyond its primary function of transporting nutrients and metabolites, the phloem serves as a conduit for hormonal and signaling molecules that regulate plant development and symbiotic efficiency. For instance, autoregulation of nodulation (AON) relies on the movement of root-derived CLE peptides via the phloem to the shoot, where they modulate *miR2111* expression to fine-tune nodule formation (Zhang et al., 2021). Additionally shoot-derived Cytokinins systemically regulate root nodulation in AON (Sasaki et al., 2014).

The phloem also mediates the translocation of critical secondary metabolites such as terpenoids, amino acids, and sucrose, which contribute to root growth and nodule maintenance under nutrient-limited conditions (Ali et al., 2021; Parsons et al., 1993). Recent three-dimensional imaging studies highlight the complexity of the vascular network within nodules, reinforcing the importance of phloem in maintaining efficient nutrient and signaling molecule exchange (Livingston et al., 2019). Despite the phloem’s critical role, the molecular basis of its regulatory function in nodulation remains inadequately characterized. While TRAP-seq has been instrumental in elucidating tissue-specific translational mechanisms, prior research has focused on cortical tissues (Song et al., 2022). The translational landscape of the phloem during nodulation remains unexplored. To address this knowledge gap, we employed a phloem-specific TRAP-seq approach to delineate the regulatory pathways governing nodulation in soybean.

Legume nodulation is a highly coordinated process involving intricate molecular and cellular interactions between host plants and nitrogen-fixing bacteria. The establishment of symplastic continuity between sieve elements and developing nodule primordia is crucial for the translocation of macromolecules, including signaling molecules and transcription factors. Previous research indicated that during the early stages of nodule development in *Medicago truncatula*, the initiation of nodule primordia induces symplastic continuity between the root phloem and the nodule, facilitating the movement of essential signaling molecules (Complainville et al., 2003). Additionally, the role of callose in regulating symplastic communication has been highlighted, as it coordinates the development of symbiotic root nodules by establishing connections between nodule primordia initials (Gaudioso-Pedraza et al., 2018). Thus, symplastic continuity not only supports physical connection but also plays a pivotal role in the signaling pathways necessary for successful nodule formation and function. The regulation of cell wall permeability and plasticity is largely governed by cell wall-modifying enzymes, including pectin methylesterase (PME) and its inhibitor (PMEI). These enzymes modulate plasmodesmata formation and maintenance, thereby influencing regulatory molecule delivery to developing nodules (Lough and Lucas, 2006).

Transcription factors expressed in the phloem orchestrate long-distance signaling by regulating hormone homeostasis and nutrient transport. Their role in coordinating rhizobial infection and nodule formation with whole-plant physiological requirements is crucial. Several families of TFs, including *Ethylene Response Factor* (*ERF*), *MYB*, *NAC*, *basic Helix-Loop-Helix* (*bHLH*), and *Auxin Response Factor* (*ARF*), have been implicated in various stages of nodulation, from rhizobial infection to mature nodule function. Early studies demonstrated that *ERF* TFs are integral to nodulation by mediating ethylene signaling, a key regulator of nodule number and function. Specifically, *ERN1* (*ETHYLENE RESPONSE FACTOR REQUIRED FOR NODULATION 1*) was shown to be a key positive regulator of Nod factor signaling in *Medicago truncatula* (Kawaharada et al., 2017). Similarly, *MYB* TFs have been linked to nodulation through their involvement in root hair infection and meristem formation, as seen in *LjMYB14*, which regulates flavonoid biosynthesis essential for rhizobial interaction in *Lotus japonicus* (Shelton et al., 2012). More recent research demonstrated that *GmMYB176* regulates isoflavonoid synthesis in soybean nodulation, emphasizing the conservation of *MYB* function in different legumes (Anguraj Vadivel et al., 2021). *NAC* TFs have also emerged as key players in the nodulation process, primarily through their role in stress responses and symbiotic regulation. For instance, *MtNAC969* was shown to enhance nodule development by modulating stress responses in *M. truncatula* (De Zélicourt et al. 2012). Additionally, *GmNAC181* in soybean was identified as a positive regulator of nodulation, highlighting a conserved role of *NAC* TFs in multiple legume species (Wang et al., 2022b).

Conversely, *ARF* transcription factors, which mediate auxin-responsive gene expression, have been implicated in the regulation of nodulation. For instance, recent studies demonstrated that expression of *GmARF8a* and *GmARF8b* is tightly controlled by *miR167c*, and this expression is essential for optimal nodule formation (Wang et al., 2015). The *bHLH* family has been implicated in root architecture modification, influencing lateral root development and nodule positioning. Studies on *MtbHLH1* indicate its role in regulating root hair growth and rhizobial attachment (Godiard et al., 2011). Furthermore, *GmbHLH176* was demonstrated to regulate iron homeostasis in soybean nodules, indicating an additional layer of metabolic regulation by *bHLH* TFs (Wu et al., 2023). While the roles of these TFs in nodulation are well known, their specific contributions within the phloem in coordinating systemic and localized nodule development remain unclear. Our study reveals dynamic changes in the expression of *ERF*, *MYB*, *NAC*, *bHLH*, and *ARF* transcription factors in the phloem during early and late nodulation stages, suggesting that phloem-mediated regulation plays a crucial role in integrating long-distance signaling with nodule formation. Collectively, our findings emphasize the importance of phloem-mediated transcriptional regulation in optimizing nodule development, linking systemic regulatory networks with root symbiosis.

## Results

### Development and Validation of a Phloem-Specific TRAP-Seq System

The TRAP-seq technique enables precise analysis of tissue-specific translatomes by leveraging a His-FLAG-tagged ribosomal protein L18 (HF-GmRPL18). This system facilitates the immunoprecipitation of actively translating polysomes and their associated mRNAs, providing a robust method for studying translation at a tissue-specific level(Zanetti et al. 2005). To establish a phloem-specific TRAP-seq system, we first identified a suitable phloem-specific promoter using laser capture microdissection (LCM) coupled with RNA-seq. Phloem tissues were isolated from soybean roots and analysis of this LCM RNA-seq dataset identified the *Glyma.01G040700* promoter as phloem-specific. Published single-cell RNA-seq data (Cervantes-Pérez et al., 2024) further validated its strong and selective expression in phloem cells, supporting its use for phloem-targeted applications (Figure 1b). Despite its high specificity, the intrinsic activity of the *Glyma.01G040700* promoter was insufficient to achieve the expression levels necessary for efficient translatome capture (Figure 1d). Robust promoter activity was essential to enhance the sensitivity of ribosome-associated mRNA isolation and ensure adequate transcript recovery for downstream analyses.

**Figure1.**
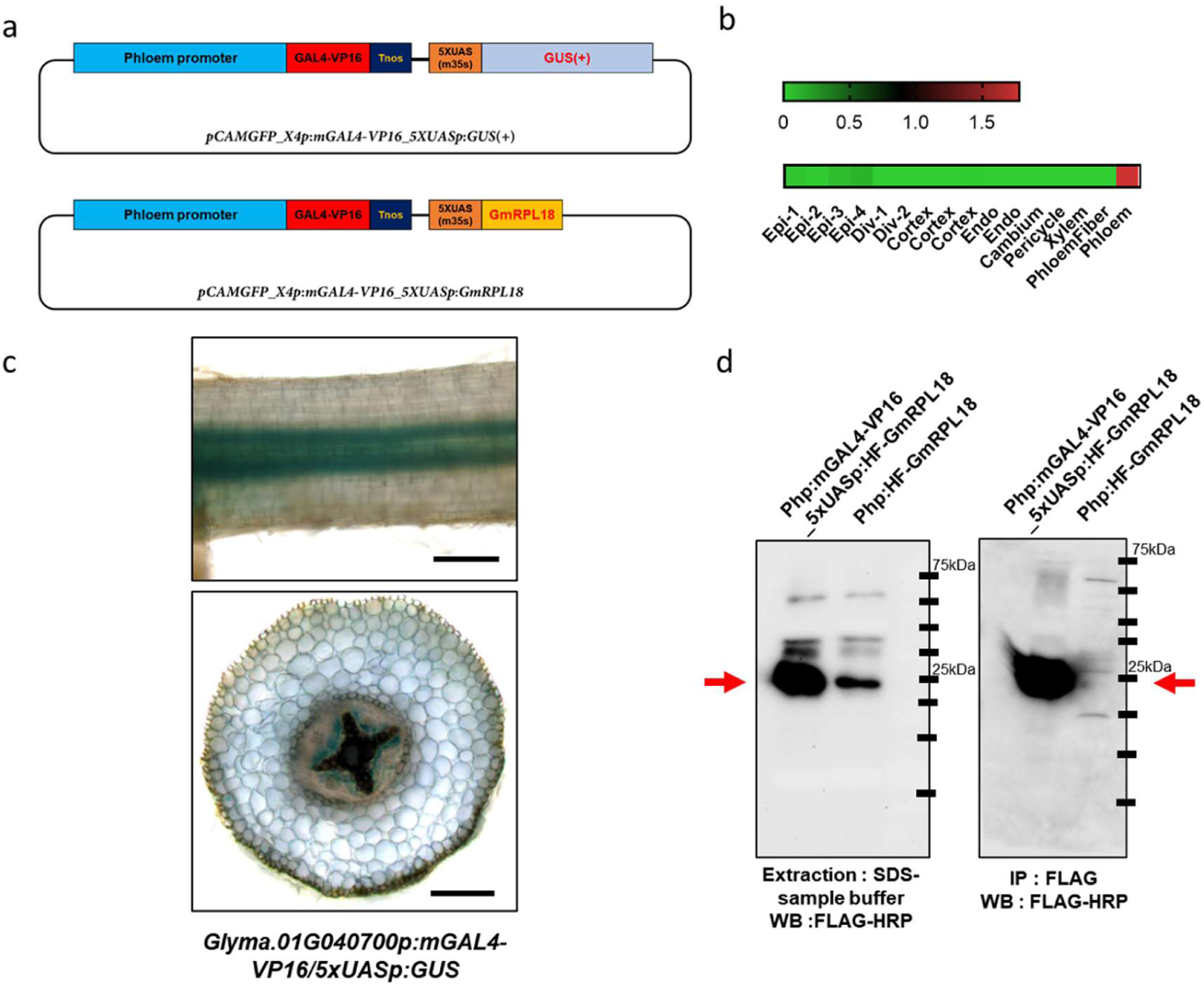
Phloem-specific TRAP-seq system design and experimental validation. (a) Schematic representation of TRAP-seq constructs. A phloem-specific promoter drives the expression of *GAL4-VP16*, which activates *UAS*-controlled *GUS* or *HF-GmRPL18* for promoter validation and ribosome tagging, respectively. (b) *In silico* single-cell RNA-seq data illustrating phloem-specific expression of *Glyma.01G040700*, confirming promoter suitability. (c) GUS reporter assay validating promoter activity. Strong phloem-specific GUS signals are observed in root longitudinal sections (top) and transverse sections (bottom). Scale bar = 0.2 mm. (d) Western blot analysis confirming HF-GmRPL18 protein expression (left, red arrow) and successful immunoprecipitation of ribosome complexes (right, red arrow), validating the functionality of the TRAP system in phloem tissue.

To overcome this limitation, we implemented the GAL4-UAS system, which significantly amplified promoter-driven expression while preserving tissue specificity (Figure 1a). In this system, *GAL4-VP16* was placed under the control of the *Glyma.01G040700* promoter, leading to a substantial increase in HF-GmRPL18 expression, as confirmed by immunoblot analysis in transgenic roots. This optimized system provided enhanced translatome capture efficiency, enabling detailed studies of phloem-specific gene regulation. To validate system functionality, we performed GUS reporter assays, which confirmed phloem-specific promoter activity (Figure 1c). Additionally, time-course TRAP-seq experiments were conducted to analyze dynamic gene expression changes in the phloem during nodulation. Immunoblot analysis of HF-GmRPL18-expressing transgenic roots demonstrated sufficient protein accumulation for efficient immunoprecipitation, validating the system’s capacity to selectively capture ribosome-associated mRNAs with high specificity and reliability (Figure 1d).

### Time-Dependent Translational Reprogramming in Soybean Root Phloem

This study aimed to investigate the translational dynamics in soybean root phloem during early (72 hpi) and late (21 dpi) stages of nodulation in response to rhizobacterial infection. The extent of translational reprogramming, visualized in the volcano plot (Figure 2a), was determined by identifying 2,636 DEGs at 72 hpi and 8,422 DEGs at 21 dpi under the criteria of FDR < 0.01 and |log2-fold change| > 1. A Venn diagram analysis (Figure 2b, Supplementary Table 1) quantified the shifts in translational activity across nodulation stages. At 72 hpi, 1,171 and 157 DEGs were uniquely upregulated and downregulated, respectively, while at 21 dpi, 3,711 and 3,403 were upregulated and downregulated.Among these, 122 DEGs transitioned from downregulated at 72 hpi to upregulated at 21 dpi. Additionally, 859 DEGs were consistently upregulated at both time points, 298 DEGs shifted from upregulated at 72 hpi to downregulated at 21 dpi, and 29 DEGs remained downregulated throughout (Figure 2b, Supplementary Table 2). These findings highlight dynamic translational regulation in soybean root phloem during nodulation. A heatmap of normalized TRAP-seq data illustrated the extent of induction for the top five upregulated DEGs at each time point, demonstrating their significant expression in rhizobium-inoculated samples (Figure 2c). RT-PCR was subsequently performed to confirm these findings, showing strong agreement with TRAP-seq data (Figure 2d). The consistency between RT-PCR and TRAP-seq results reinforces the reliability of translational profiling in capturing stage-specific regulatory changes.

**Figure 2.**
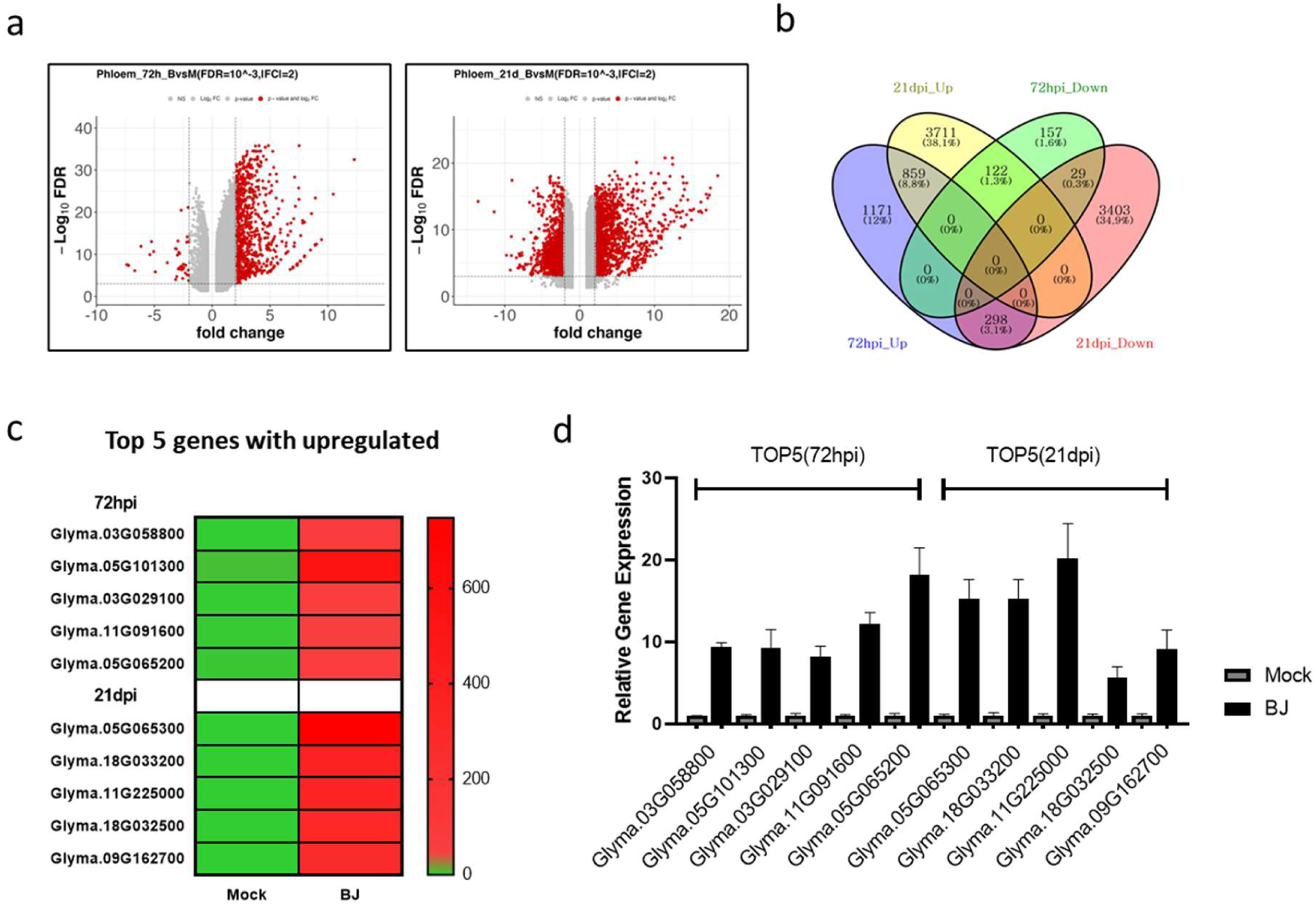
Differential Gene Expression Analysis in Soybean Root Phloem Using TRAP-Seq and Validation. (a) Volcano plots of DEGs in root phloem at 72 hpi (left) and 21 dpi (right) with mock and rhizobium inoculation. Significant genes (FDR < 0.1, |log2 FC| ≥ 2) are shown in red. (b) Venn diagram showing overlap of upregulated and downregulated genes between 72 hpi and 21 dpi conditions. (c) Heatmap of normalized RNA-seq expression values for the top 5 upregulated genes at 72 hpi and 21 dpi. (d) Bar graph showing Real-time PCR validation of the top 5 upregulated genes, displaying relative expression levels compared to mock conditions.

### Functional Genomics and Metabolic Pathways Underlying Root Phloem’s Role in Nodulation

To elucidate the translational regulatory mechanisms governing phloem function during nodulation, we conducted KEGG pathway and GO enrichment analyses to identify key biological processes enriched in phloem-associated transcripts at 72 hpi and 21 dpi, using REVIGO to summarize and reduce redundancy in GO terms for clearer interpretation (Supplementary Table 3, 4).

At 72 hpi, upregulated pathways fall into three main categories: (1) structural and defense responses, including cell wall biogenesis and ethylene signaling, which facilitate early nodulation-related adaptations and the formation of a symplastic network; (2) hormonal and regulatory pathways, such as jasmonic acid and brassinosteroid signaling, that coordinate nodulation responses and cell communication; and (3) transport functions, including phloem transport and amino acid regulation, establishing initial nutrient mobilization essential for nodule development (Figure 3a, Figure S1a). By 21 dpi, enriched pathways shift to (1) transport-related processes, such as phosphate ion homeostasis, reflecting the phloem’s role in regulating nutrient movement into developing nodules; (2) structural modifications, including vascular differentiation and cell wall modification, reinforcing nodule vasculature for sustained transport; and (3) regulatory mechanisms like amino acid transport and mechanical stimulus response, ensuring proper coordination of mature nodule function (Figure 3c, Figure S1c). Concurrently, downregulated pathways at 21 dpi indicate a shift from early proliferation to functional specialization, categorized as (1) reduced carbohydrate metabolism, including starch and sucrose metabolism, prioritizing nitrogen assimilation; (2) decreased cell proliferation, marked by lower nuclear division and mitotic activity, indicating meristem transition; and (3) suppressed biochemical pathways, such as glycosyl metabolism and DNA recombination, suggesting stable genomic regulation in mature nodules (Figure 3d, Figure S1c). These findings highlight the role of the phloem in first establishing and later regulating the symplastic transport network essential for effective nodule function and nitrogen fixation (Complainville et al., 2003).

**Figure 3.**
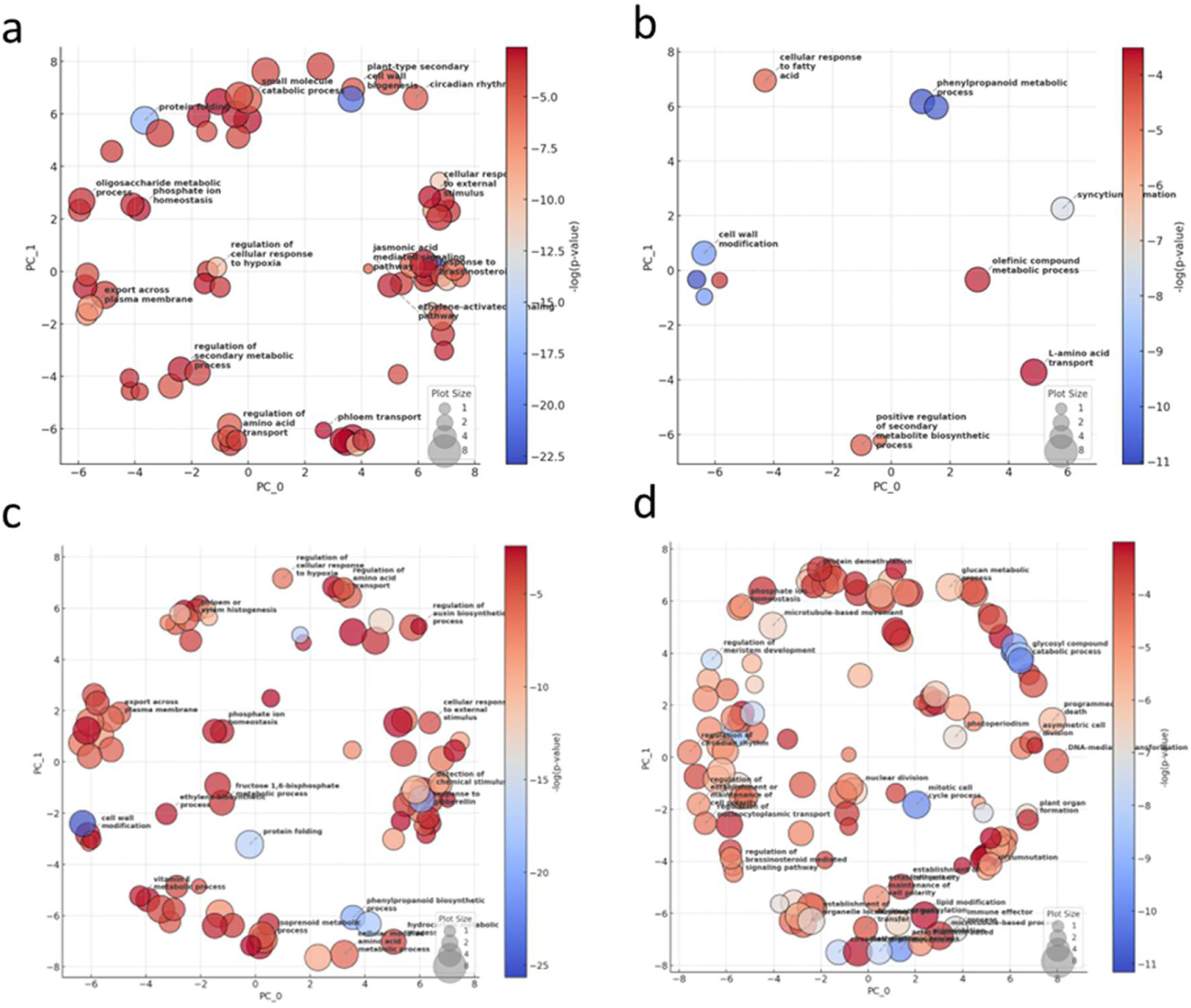
GO Enrichment Analysis of Differentially Expressed Genes in Soybean Root Phloem. REVIGO visualization of biological process GO enrichment based on DEGs at 72 hpi and 21 dpi. Each sphere represents a GO term, positioned by semantic similarity, with color indicating -log10(P-value) and size representing term frequency. (a) Upregulated genes at 72 hpi, (b) Downregulated genes at 72 hpi, (c) Upregulated genes at 21 dpi, (d) Downregulated genes at 21 dpi.

### Temporal and Spatial Expression Patterns of PME and PMEI Genes in Soybean Root Nodulation

The symplastic transport network plays a fundamental role in nodule formation and function. Given the observed regulation of various cell wall-modifying enzymes by bacterial infection in the phloem, we conducted a detailed analysis of the expression dynamics of *PME* and its inhibitor (*PMEI*) to elucidate their involvement in plasmodesmata regulation. These enzymes are critical for maintaining cell wall permeability and modulating symplastic connectivity, essential processes for coordinated intercellular communication during nodulation.

Phloem-TRAP-seq analysis and genome-wide studies revealed distinct temporal and spatial expression patterns of *PME* and *PMEI* genes during soybean nodulation, categorizing them into four functional groups (Wang et al., 2020; Wang et al., 2021). Group I *PME* genes, significantly upregulated at 72 hpi and maintained at 21 dpi, largely overlap with PMEI Group I genes, both containing a PMEI domain that enables self-regulation of PME activity. This regulation balances cell wall flexibility and reinforcement, potentially contributing to plasmodesmata remodeling and the establishment of symplastic connectivity during nodulation (Figure 4a, b, c). In contrast, several Group III and IV *PME* genes were predominantly expressed at later stages, contributing to nodule maturation and structural stabilization (Figure 4a) (Wang et al., 2021). The upregulation of both *PMEI* Group I and Group III genes at 72 hpi and 21 dpi, with most containing PME domains, suggests that self-regulation is essential for coordinated cell wall modifications (Figure 4a, c). These findings indicate that *PME* and *PMEI* genes not only mediate cell wall modifications but also regulate plasmodesmata connectivity, ultimately influencing nodulation efficiency.

**Figure 4.**
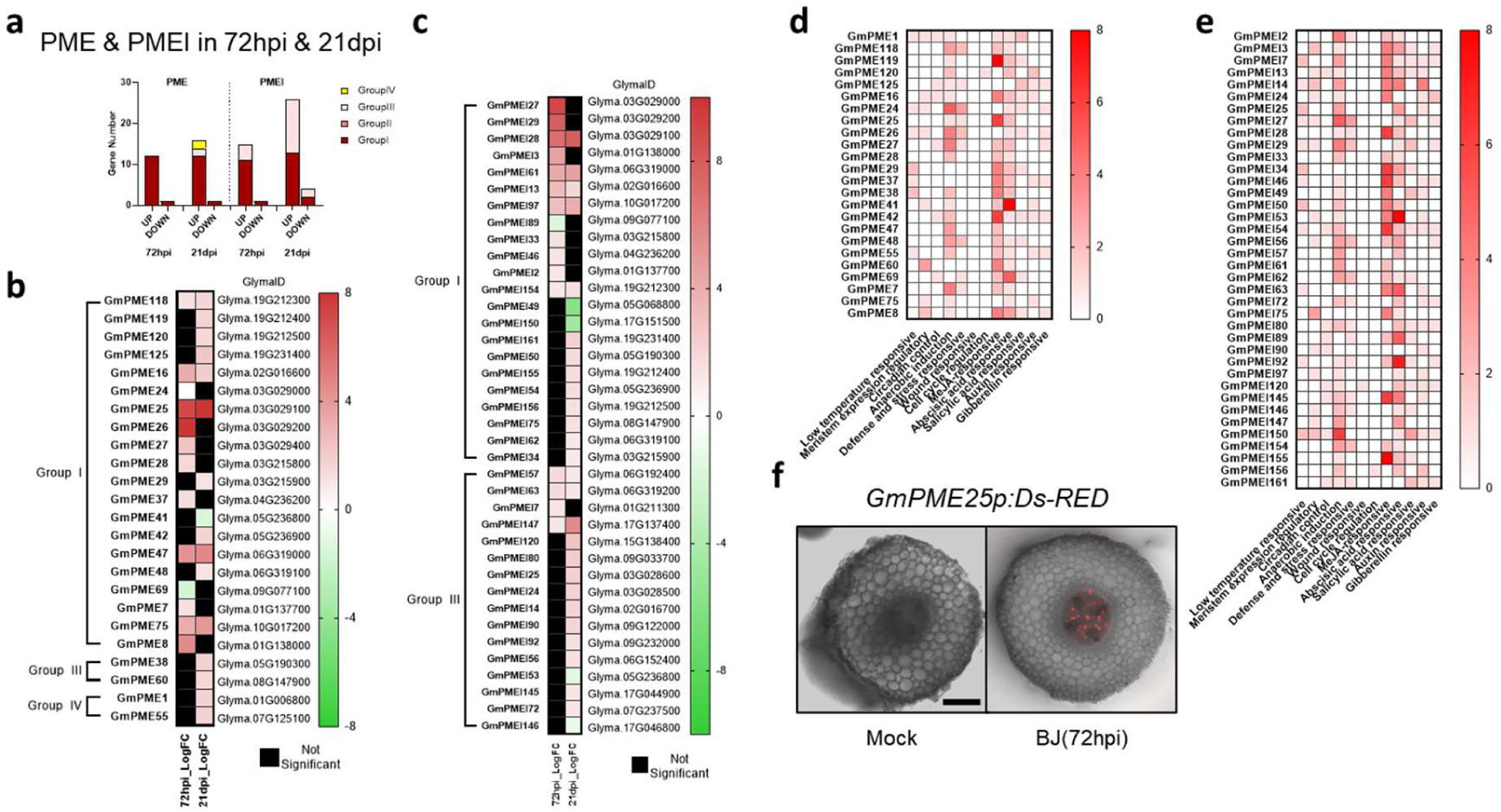
Temporal Expression and Regulatory Analysis of PME and PMEI Genes in Soybean Root Phloem During Nodulation. (a) Classification and distribution of differentially expressed *PME* and *PMEI* genes at 72 hpi and 21 dpi. (b, c) Heatmaps depicting log2 fold change (log2FC) in PME (b) and PMEI (c) gene expression over time. (d, e) Cis-regulatory element analysis of PME (d) and PMEI (e) promoters, with color intensity indicating element frequency. (f) Ds-RED fluorescence assay showing *GmPME25* promoter activity in soybean root phloem at 72 hpi. Scale bar = 0.2 mm.

Cis-element analysis of *PME* and *PMEI* promoter regions using PlantCARE identified MeJA-responsive, ABA-responsive, and anaerobic induction-related elements in both *PME* and *PMEI* genes (Figure 4d, e). These findings suggest that *PME* and *PMEI* expression is modulated by jasmonic acid, abscisic acid, and oxygen availability, aligning their activity with environmental and hormonal cues during nodulation. Functional validation using *GmPME25p:Ds-RED* confirmed *GmPME25* activation in root phloem at 72 hpi, reinforcing its role in early nodule development (Figure 4f). However, RNAi-mediated suppression of Group I *PME* genes did not yield significant phenotypic alterations, suggesting potential functional redundancy within the *PME* gene family (data not shown). Collectively, these findings provide a comprehensive framework for understanding the regulatory mechanisms governing *PME* and *PMEI* expression during nodulation.

### Dynamic Regulation of Transcription Factors in Soybean Root Phloem During Nodulation

Transcription factor (TF) control plays a crucial role in the symbiosis between legumes and rhizobia. However, the temporal roles of major TFs in the phloem remain unclear despite its essential function in systemic signaling. This study analyzed TF expression in the root phloem at 72 hpi and 21 dpi, revealing stage-specific regulatory shifts. At 72 hpi, *ERF* (gene ID: 34) exhibited the highest increase in expression, followed by upregulation of *C2H2* (gene ID: 13), *NAC* (gene ID: 26), *MYB* (gene ID: 17), *WRKY* (gene ID: 21), *bHLH* (gene ID: 16), and *LBD* (gene ID: 10) (Figure 5a). Minimal reductions in expression were observed, with only slight suppression of ERF (2) (Figure 5a). These findings highlight stage-specific TF regulation during nodulation. In the early phase, ERF plays a central role in ethylene signaling, while *C2H2* and *NAC* contribute to early stress response and nodule development. *MYB* supports root hair infection and flavonoid biosynthesis, *WRKY* is involved in metabolic defense regulation, and *bHLH* influences root architecture and rhizobial attachment. At 21 dpi, *ERF* (gene ID: 49) and *MYB* (gene ID: 53) showed the highest expression levels, followed by consistent upregulation of *bHLH* (gene ID: 35) and *NAC* (gene ID: 30). *C2H2* (gene ID: 21) maintained stable expression, while *ARF* (gene ID: 10) exhibited the most significant reduction. *WRKY* (gene ID: 19) and *C2H2* (gene ID: 12) were downregulated, and moderate suppression was observed in *bHLH* (gene ID: 20) and *MYB* (gene ID: 12) (Figure 5a). In the maturation phase, *MYB* and *ERF* persist as central regulators, facilitating secondary metabolism and vascular differentiation, and ensuring the developmental progression of nodules. The decline in *ARF* expression at 21 dpi suggests a regulatory mechanism that negatively regulates nodulation, whereas the reduced expression of *WRKY* and *C2H2* indicates a shift away from early-stage defense activation.

**Figure 5.**
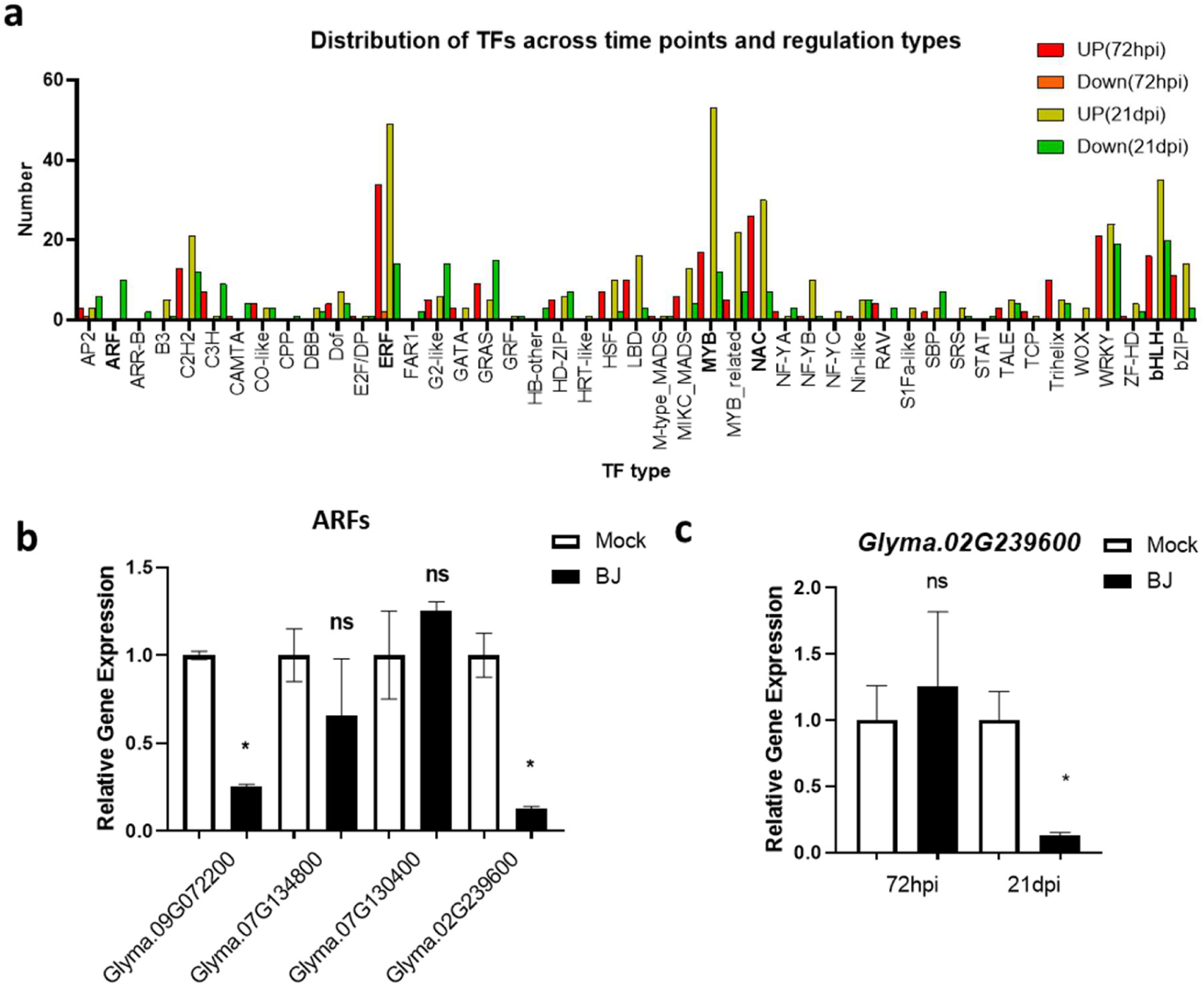
Transcriptional Regulation of TFs During Nodulation. (a) Dynamic expression patterns of transcription factors (TFs) at 72 hpi and 21 dpi, highlighting key TF families including *ARF*, *NAC*, *MYB*, *ERF*, and *bHLH*. (b) qRT-PCR validation showing significant downregulation of *ARF* genes at 21 dpi. (c) Time-course expression analysis of *Glyma.02G239600*, illustrating its stage-specific transcriptional regulation during nodulation.

The coordinated regulation of these TFs ensures systemic signaling integration and optimized nodulation efficiency in the phloem. To comprehensively validate the stage-specific regulation of *ARF* genes, we conducted quantitative Real-time PCR analysis on four *ARF* genes at 21 dpi

(Figure 5b). The analysis revealed that *Glyma.09G072200* and *Glyma.02G239600* exhibited significant transcriptional repression in the 21 dpi TRAP-seq dataset, confirming their downregulation during late-stage nodulation. Furthermore, a time-course expression analysis of *Glyma.02G239600* (Figure 5c) demonstrated a clear decline in expression, providing additional evidence of its temporal regulation. These findings suggest that ARF suppression in the phloem may play a role in regulating auxin signaling at 21 dpi, potentially contributing to optimal nodulation.

### Identification of *GmbHLH121* as a Key Nodulation Regulator

One of the basic Helix-Loop-Helix (bHLH) transcription factors, encoded by *Glyma.03G105700*, a Myc-like *GmbHLH121*, was identified as a strong candidate for nodulation regulation. This identification was based on its enrichment in the phloem at 72 hpi and 21 dpi. The high expression pattern observed in root hair-stripped samples following rhizobacterial inoculation further supports its role in nodulation (Figure S2). Promoter activity analysis using GUS reporter assays demonstrated phloem-specific induction of the *GmbHLH121* promoter in response to rhizobacterial infection. This finding was consistent with phloem-specific responses observed in TRAP-seq data (Figure 6a). To functionally characterize *GmbHLH121*, hairy root transformation was performed to generate transgenic lines with either RNAi-mediated knockdown (RNAi) or overexpression (OX). Phenotypic analysis revealed that RNAi lines exhibited increased nodule numbers, while OX lines did not differ in nodulation, demonstrating the TF’s critical role in phloem-specific nodule formation (Figure 6b). Real-time PCR confirmed successful genetic modifications, with decreased *GmbHLH121* expression in RNAi lines and increased expression in OX lines (Figure 6c).

**Figure 6.**
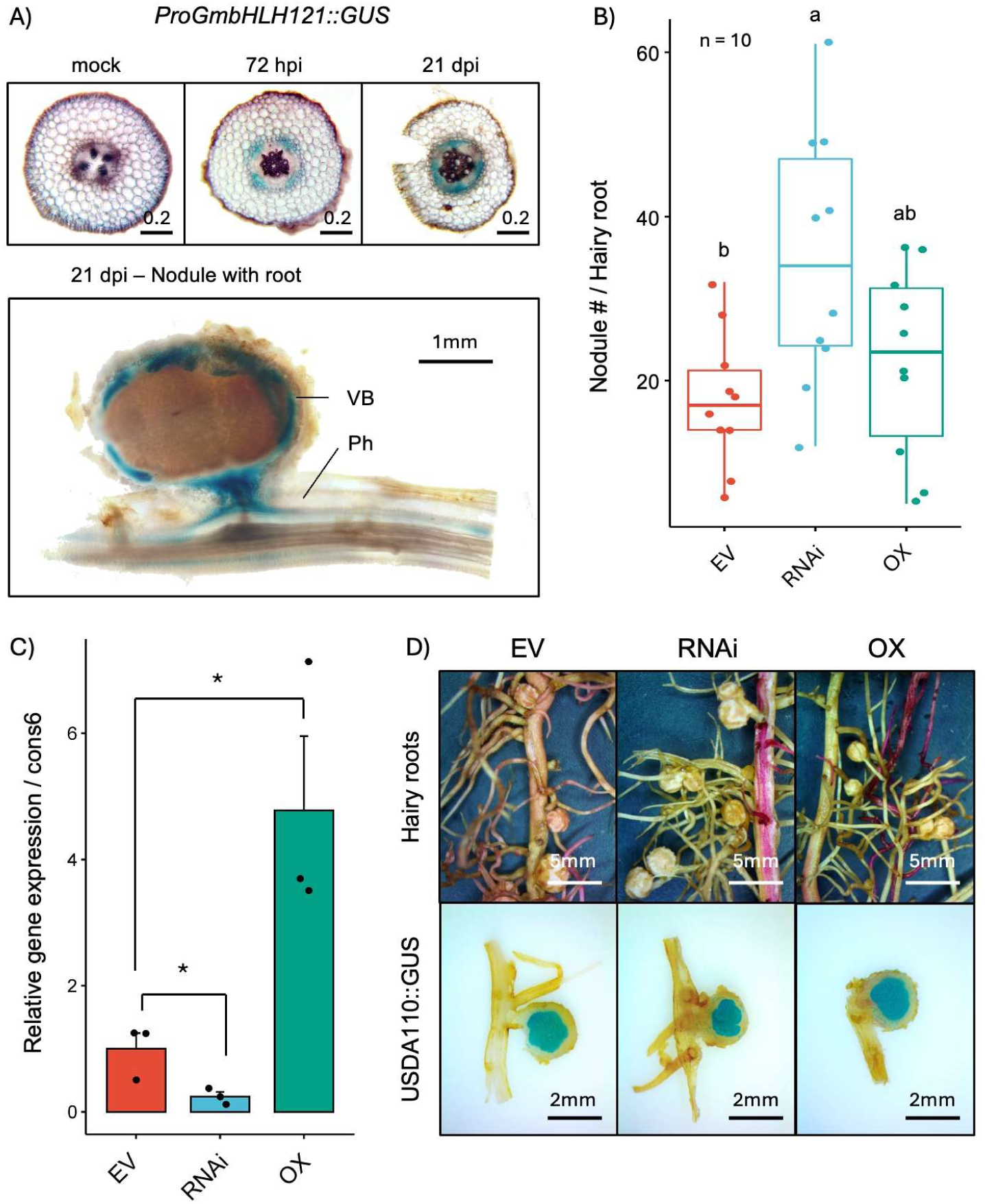
Expression and Functional Characterization of *GmbHLH121* in Soybean Nodulation. Histochemical, phenotypic, and molecular analyses of *GmbHLH121* expression in soybean roots and nodules. (a) Cross-sections of root tissues displaying *GmbHLH121* promoter-driven GUS activity at different developmental stages, highlighting expression in vascular tissues and nodules. (b) Nodule phenotypic analysis of *GmbHLH121* RNAi and overexpression (OX) lines, illustrating morphological differences in nodule development (n = 10, statistical significance denoted by letters). (c) Bar graph depicting relative *GmbHLH121* expression levels in RNAi and OX lines, quantified via qRT-PCR using *cons6* as the control, with asterisks indicating statistically significant differences. (d) Hairy root phenotypic analysis comparing Empty Vector, RNAi, and OX lines, accompanied by nodule structural assessment using *USDA110::GUS* staining to visualize bacteroid. Whole-mount images of nodulated roots and corresponding cross-sections confirm spatial expression patterns. Scale bars: 0.2 mm (a, top), 1 mm (a, bottom), 5 mm (d, top), 2 mm (d, bottom).

Histological analysis indicated that nodule structures remained consistent across all transgenic lines, suggesting that *GmbHLH121* primarily regulates nodule number rather than morphology (Figure 6d). Interestingly, the expression of early-stage nodulation markers, such as *ENOD40*, *NIN*, and *NSP*, did not differ among transgenic plants (Figure S3). This suggests that the pathway controlling nodule numbers might operate independently of these genes or function downstream.

## Discussion

### Establishment and Functional Validation of a Phloem-Specific TRAP-Seq System

The establishment of a phloem-specific TRAP-seq system represents a significant advancement in elucidating translational regulatory mechanisms in the phloem during nodulation. By utilizing the *Glyma.01G040700* promoter, we successfully targeted ribosome-associated mRNAs in phloem tissues. However, initial assessments revealed that the endogenous activity of this promoter alone was insufficient to achieve optimal ribosome-tagging efficiency for TRAP-seq. To address this limitation, we introduced the GAL4-UAS system, which significantly enhanced expression levels while maintaining phloem specificity. The successful implementation of the GAL4-UAS amplification strategy underscores the necessity of optimizing promoter-driven expression in tissue-specific TRAP-seq applications. While our previous study effectively utilized endogenous promoters for TRAP-seq in cortex tissue (Song et al., 2022), our findings suggest that additional regulatory elements may be required for applications targeting the phloem in soybean. One of the key advantages of this optimized system is its ability to capture dynamic translational changes in the phloem during nodulation. Time-course TRAP-seq experiments provided high-resolution insights into phloem translatome reprogramming, facilitating the identification of transcription factors and signaling molecules crucial for phloem regulation. Furthermore, this system minimizes contamination from adjacent cell types, thereby increasing the reliability of detected gene expression changes and enhancing the accuracy of subsequent functional analyses. Despite these advances, several limitations must be considered. The translational changes observed in the phloem do not necessarily indicate phloem-specific regulation, as similar changes may occur in neighboring root tissues. A more comprehensive comparative approach incorporating TRAP-seq datasets from multiple root tissues is required to rigorously delineate phloem-specific translational regulation.

While our previous research included similar experiments in cortex tissues, data from additional root tissues remains insufficient. To improve the specificity of phloem-regulation, integration with single-cell RNA-seq analysis is essential. Although single-cell RNA-seq data for nodules are available (Liu et al., 2023; Cervantes-Pérez et al., 2024), root-specific single-cell RNA-seq data comparing pre- and post-rhizobacterial infection states are still lacking. This data gap constrains our ability to precisely map phloem-specific expression changes. Future studies should prioritize generating these datasets and developing computational strategies to enhance the resolution of tissue-specific gene expression analysis. Nonetheless, in the absence of such datasets, our TRAP-seq analysis provides a crucial framework for understanding the functional changes associated with nodulation specifically within the phloem. By offering novel insights into phloem-specific translational regulation, our study establishes a foundational platform that can be leveraged for future investigations into legume symbiosis and phloem-mediated regulatory networks.

### Phloem-Driven Immune Regulation in Legume-Rhizobium Interactions

Rhizobium infection initiates in root hairs and spreads to the cortex, during which the *Nod factor* (*NF*) plays a crucial role in modulating innate immune signaling through interactions with the host’s *pattern recognition receptor*s (*PRR*s) to facilitate symbiosis. Generally, *PRR*s recognize *microbial-associated molecular patterns* (*MAMP*s) to activate defense responses, but the Nod factor secreted by Rhizobium partially suppresses *PRR* signaling, thereby modulating the host immune response and enabling discrimination between pathogenic and symbiotic microbes (Liang et al., 2013; Cao et al., 2017). Following initial infection in root hairs, as it progresses into the cortex, the immune response remains finely regulated to facilitate Rhizobium establishment, with genes such as *RIN4*, *WRKY22*, and *CDPK*s involved in sophisticated mechanisms that distinguish pathogens from Rhizobium (Yu et al., 2018; Feng et al., 2021; Wang et al., 2022a; Tóth et al., 2023). This regulatory mechanism is essential for maintaining the balance between symbiosis and immunity.

Casparian strips generally function as selective barriers within endodermal cells, regulating water and nutrient uptake while restricting pathogen invasion (Robbins et al., 2014). While apoplastic pathway-based pathogen infection is effectively blocked by Casparian strips, the potential for infection through the symplastic pathway remains primarily facilitated by plasmodesmata. Our study provides strong evidence that a sustained increase in the expression of various cell wall-modifying enzymes in the phloem from early to late infection stages strongly correlates with plasmodesmata regulation, suggesting that Rhizobium infection may induce structural modifications that enhance plasmodesmata permeability, thereby facilitating increased molecular transport, particularly macromolecule movement, between the phloem and nodule. Previous studies reported increased macromolecule transport from the phloem to nodule primordia following Rhizobium infection(Complainville et al., 2003), further supporting the hypothesis that plasmodesmata opening between the phloem and cortex is a key regulatory mechanism in both infection and symbiosis.

In our study, we confirmed a significant increase in the expression of *GmBAK1* (*Glyma.05G119600*), *GmWRKY22* (*Glyma.16G031900*), *GmLRRs* (*Glyma.09G210600*, *Glyma.01G010700*, *Glyma.08G128800*, *Glyma.18G093900*), and GmCDPKs (*Glyma.10G084000*, *Glyma.02G192700*) in the phloem at 72 hpi, supporting the hypothesis that the phloem serves as a key immune regulatory hub to control infection spread. Unlike the fine-tuned immune gene regulation observed in the cortex and root hairs, the phloem displayed a more pronounced upregulation of multiple immune-related genes. This suggests that beyond immune modulation, the phloem plays a proactive role in restricting pathogen spread. Furthermore, the accumulation of immune-regulatory proteins in the phloem may contribute to both pathogen resistance and symbiotic signaling, potentially facilitating local immune responses and coordinating signal transduction from the phloem to the cortex.

Thus, our findings suggest that the phloem functions not only as a transport conduit but also as a key regulatory tissue that precisely modulates infection spread through immune responses and molecular exchange during Rhizobium infection. Further studies are necessary to explore the interplay between plasmodesmata regulation and immune networks within the phloem at a more mechanistic level.

### Regulation of AAP2 and AAP8 Expression and Their Role in Nitrogen Fixation in Root Nodules

In legume root nodules, bacteroids fix atmospheric nitrogen (N₂), converting it into ammonia (NH₃), which is then transported into host cells via the Nod26 channel and subsequently assimilated into glutamine and asparagine (Fortin et al., 1987). The transport of fixed nitrogen varies across species; temperate legumes (e.g., pea and alfalfa) primarily utilize amide forms, whereas tropical legumes transport nitrogen as ureides, such as allantoin and allantoic acid (Miao et al., 1991; Atkins and Smith, 2000). These organic nitrogen compounds are exported from nodules into the xylem and distributed throughout the plant via interactions with the phloem (Herridge et al., 1978)

The endodermis of root nodules restricts apoplastic nitrogen transport, necessitating symplastic movement. In soybean, UPS1-1 and UPS1-2 transporters play crucial roles in moving allantoin and allantoic acid (Pélissier et al., 2004; Collier and Tegeder, 2012). Meanwhile, in Arabidopsis, AAP2 and AAP6 mediate amino acid exchange between the xylem and phloem, facilitating efficient nitrogen redistribution (Hunt et al., 2010; Zhang et al., 2010). AAP8 also functions as a key regulator of source-to-sink nitrogen partitioning, influencing phloem nitrogen transport and seed development (Santiago and Tegeder, 2016). Additionally, in legumes, AAP and UPS1 transporters are localized to xylem parenchyma and transport phloem, suggesting their role in transient nitrogen storage and organic nitrogen transfer (Pélissier et al., 2004; Tegeder et al., 2007).

Previous studies identified enrichment of the gene ontology (GO) term “regulation of amino acid transport” at 72hpi and 21dpi (Figure 3a, c). A detailed examination of *AAP* gene expression in phloem revealed that *GmAAP2* (*Glyma.11G107000*, *Glyma.06G156700*) expression is significantly upregulated at 72 hpi in phloem. Given that AAP2 is a major transporter responsible for transferring amino acids from the xylem to the phloem, this may play a role in providing amino acids and energy sources required for cell proliferation and expansion during early nodule formation. However, by 21dpi, *GmAAP2* (*Glyma.11G107000*, *Glyma.12G032000*) expression showed a declining trend, likely due to reduced dependency on external amino acid supply as nodules mature. This decrease may also be attributed to the ability of bacteroids to autonomously fix nitrogen, reducing the necessity for phloem-mediated nitrogen input. These findings suggest that *AAP2* plays a pivotal role in regulating early nitrogen supply, while its function gradually diminishes as nodules mature.

*AAP8* is a major regulator of amino acid loading into the phloem, potentially facilitating the efficient distribution of nitrogen fixation products to the shoot and seeds in Arabidopsis. Our findings indicate that *GmAAP8* expression remained unchanged at 72hpi but significantly increased at 21dpi. This suggests that as nodules mature and nitrogen fixation becomes active, the demand for redistribution of glutamine, asparagine, and ureides (allantoin, allantoic acid) to sink tissues (e.g., seeds and emerging leaves) increases. Therefore, the elevated expression of *GmAAP8* in the phloem could potentially serve as a key regulatory mechanism for the efficient partitioning of fixed nitrogen.

Our study suggests that in soybean, *GmAAP2* is primarily responsible for early nitrogen supply, whereas *GmAAP8* may contribute to the later-stage distribution of nitrogen fixation products. *GmAAP2* may be involved in nitrogen acquisition and support nodule growth during the early stages, whereas *GmAAP8* enhances nitrogen allocation to developing tissues at later stages, ultimately optimizing nitrogen use efficiency (NUE). Future research should focus on elucidating the regulatory networks governing their expression and integrating these insights into crop improvement strategies for enhanced nitrogen utilization efficiency.

### Regulation of Phloem-Associated *PME* and *PMEI* in Nodulation: Insights into Cell Wall Modification and Symplastic Transport

Nodulation in legumes is a complex developmental process that appears to require intricate coordination of symbiotic signaling, cell wall remodeling, and plasmodesmata regulation. In this study, our findings suggest that the majority of *PME* genes activated at 72 hpi and 21 dpi contain a *PMEI* domain. Previous research suggests that *PME* isoforms possessing an intrinsic *PMEI* domain may have a more refined and responsive regulatory mechanism compared to those lacking it (Bosch and Hepler, 2005). This regulation appears to be influenced by pH, as interactions between PME and PMEI are pH-dependent, leading to dynamic shifts in enzymatic activity (Hocq et al., 2021). Additionally, PMEIs play a multifaceted role beyond direct inhibition, contributing to broader regulatory networks involved in cell wall remodeling and stress responses (Wormit and Usadel 2018). This inherent regulatory capacity could be particularly advantageous in highly dynamic processes such as rhizobial infection and nodulation, where timely modifications of the cell wall might be essential for both bacterial entry and nodule organogenesis.

Cell wall modifications mediated by *PME* and *PMEI* seem to play a role in the spatial and temporal regulation of nodulation. During rhizobial infection, localized cell wall acidification and reactive oxygen species (ROS) signaling in root hair tips were reported to contribute to symbiotic establishment (Ramu, Peng, and Cook 2007). The activation of ROS signaling during infection may lead to pH fluctuations that influence *PME* activity, potentially promoting rapid and dynamic modifications of the cell wall at the site of bacterial entry. Since *PMEI* binding affinity to *PME* is pH-dependent, transient acidification of the infection site could weaken *PMEI* inhibition, allowing for localized *PME* activation and subsequent remodeling of the cell wall. This mechanism may facilitate the controlled relaxation of cell wall structures necessary for bacterial invasion, infection thread formation, and the early stages of nodule development.

In addition to its possible role in early infection, *PME* and *PMEI* gene regulation may also contribute to the long-distance coordination of nodulation. The observed increase in *PME-PMEI* gene expression in phloem-associated tissues suggests a potential role in modulating plasmodesmata permeability, thereby influencing the systemic transport of signaling molecules that are likely critical for nodulation. Studies in *Medicago truncatula* demonstrated that callose degradation enhances plasmodesmata conductivity, ultimately promoting rhizobial infection and increasing nodulation efficiency (Gaudioso-Pedraza, 2017). Given the functional parallels in plasmodesmata regulation, it is plausible that *PME-PMEI* activity in phloem tissues contributes to symplastic connectivity adjustments that optimize long-distance signaling and resource allocation during nodulation, though further research is necessary to validate this hypothesis.

Further regulatory complexity is introduced by environmental factors that influence *PME* and *PMEI* expression. Cis-element analysis of their promoter regions revealed motifs responsive to jasmonic acid (MeJA), abscisic acid (ABA), and hypoxia-inducible elements. These findings suggest that *PME* and *PMEI* genes are tightly regulated by hormonal and environmental cues, particularly in response to the fluctuating conditions encountered during nodulation. Jasmonic acid and abscisic acid, well-characterized regulators of plant defense and abiotic stress responses, may orchestrate *PME* and *PMEI* activity to balance nodulation with overall plant fitness. However, the precise regulatory mechanisms linking these signals to *PME* and *PMEI* activity remain to be fully elucidated.

To further investigate the functional role of *PME* during nodulation, transgenic soybean lines expressing *GmPME25p:Ds-RED* indicated its activation in root phloem at 72 hpi, supporting its involvement in regulating cell wall plasticity in early infection stages. However, RNAi-mediated suppression of Group I *PME* genes did not produce significant phenotypic alterations (data not shown), suggesting the existence of functional redundancy within the *PME* gene family. This redundancy likely serves as a compensatory mechanism to ensure the robustness of nodulation despite fluctuations in the expression of individual genes.

Taken together, these findings suggest that *PME* and *PMEI* may function as multifaceted regulators in soybean nodulation. Beyond their canonical role in cell wall remodeling, these enzymes may contribute to infection site plasticity, long-distance symbiotic signaling, and environmental adaptation during nodule development. Nonetheless, further research is required to clarify the precise molecular mechanisms by which *PME* and *PMEI* coordinate these processes, particularly in relation to plasmodesmata regulation and hormonal cross-talk.

### Phloem as a Regulatory Hub: Ethylene and Auxin Signaling in Nodulation

The regulatory dynamics of transcription factors within the phloem play a central role in the legume-rhizobium symbiotic interaction. This study refines previous transcriptional models by providing high-resolution insights into phloem-specific regulatory mechanisms, reinforcing its function as an active signaling hub rather than a passive transport conduit. Prior research established that *MYB* governs flavonoid biosynthesis to enhance rhizobial infection (Liu et al., 2013), *bHLH* modulates nodule vascular patterning and development in *Medicago truncatula* (Godiard et al., 2011), *C2H2* and *NAC* mediate symbiotic signaling and defense responses (De Zélicourt et al., 2012; Liu et al., 2022), and *WRKY* regulates pathogen defense and metabolic activation (Yang et al., 2024). Our findings confirm that these TFs undergo significant expression shifts within the phloem, underscoring their role as a regulatory nexus in nodulation. Notably, we highlight the dynamic expression of *ERF* and *ARF* as key mediators of phloem-driven long-distance signaling during nodulation.

*ERF* exhibited the highest upregulation during the early nodulation stage (72 hpi), suggesting that ethylene signaling through the phloem is integral to nodulation initiation. Previous studies established that *ERF* facilitates infection thread formation and orchestrates nodulation onset within the root (Kawaharada et al., 2017), while ethylene synthesized in the shoot has been proposed as a systemic regulator of root nodulation (Penmetsa and Cook, 1997). Our findings validate and extend this model by demonstrating that *ERF* undergoes significant phloem-specific upregulation, supporting the hypothesis that shoot-derived ethylene is translocated via the phloem to regulate nodulation. This suggests a sophisticated layer of control in which ethylene signaling not only serves as a transportable cue but also fine-tunes *ERF* activity to precisely modulate nodulation initiation.

Conversely, *ARF* was markedly downregulated during the late nodulation stage (21 dpi), suggesting that auxin signaling suppression through the phloem plays a crucial role in refining nodulation dynamics. Prior research has demonstrated that *miR167c*, expressed within vascular tissue, targets *GmARF6* and *GmARF8* to regulate nodule formation within the root (Yan et al., 2015; Zhang et al., 2021). In alignment with these findings, our data reveals a pronounced reduction in *ARF* expression, particularly *ARF8* (*Glyma.02G239600*, *Glyma.11G204200*), within the phloem at 21 dpi. This suggests that *miR167c*-mediated suppression of *ARFs* may extend beyond local root regulation, instead constituting a shoot-to-root long-distance signaling mechanism via the phloem. The downregulation of auxin signaling at this stage likely promotes nodule maturation while preventing excessive nodule formation, optimizing symbiotic efficiency.

Taking together, this study presents a refined model of nodulation regulation by emphasizing the role of phloem-specific transcriptional changes. The early upregulation of *ERF* in the phloem supports nodulation initiation, while the late downregulation of *ARF* fine-tunes nodule development, preventing excessive formation. By integrating high-resolution transcriptional profiling, we provide a deeper understanding of the systemic regulatory networks governing soybean nodulation, elucidating the phloem’s essential role as a conduit for long-distance signaling.

### Phloem-Localized *GmbHLH121* and Its Role in Nodule Development

*GmbHLH121*, one of the 355 homologous basic helix-loop-helix (bHLH) transcription factors (TFs) identified in *Glycine max*, is classified within subclade IVa, the most expansive among the 25 established bHLH subclades. Notably, legume species exhibit a greater diversification within this subclade than other plant families, suggesting a lineage-specific expansion and functional divergence (Brown and Hudson, 2015; Suzuki et al., 2021). Within subclade IVa, *GmbHLH121* is assigned to subgroup 1, which is characterized by a broader range of expression profiles, in contrast to subgroups 2 and 3 that predominantly exhibit nodule- and root-specific expression patterns.

Recent findings established *GmbHLH300* as a negative regulator of nodulation and nitrogen fixation, particularly under iron-deficient and iron-excess conditions, thereby playing a pivotal role in iron homeostasis within nodules (Wu et al., 2023). Mechanistically, *GmbHLH300* directly associates with the promoter regions of *ENOD93*, a positive regulator of nodulation, and *GmLba*, a gene encoding a key enzyme for leghemoglobin biosynthesis, thereby modulating their transcriptional activity.

In our investigation, *GmbHLH121* emerged as a previously uncharacterized regulatory factor in nodulation, exhibiting a significant role in the response to rhizobial inoculation and modulating nodule numbers. However, differential gene expression analyses revealed that key early nodulation marker genes exhibited no substantial variation in expression levels across empty vector (EV), overexpression, and knockdown lines. This observation suggests that *GmbHLH121* does not act as a direct transcriptional regulator of these early nodulation-associated genes, indicating a more intricate regulatory mechanism that warrants further in-depth functional characterization.

## Materials and Methods

### Plant Materials, Growth Conditions, and Bacterial Strains

Soybean (*Glycine max*) cultivar Williams 82 was used for all experiments. Seeds were surface sterilized and germinated on sterilized germination paper or in a 3:1 mixture of vermiculite and perlite under controlled conditions (16 h light/8 h dark, 26°C/23°C light/dark, 80% relative humidity). Seedlings were inoculated with *Bradyrhizobium diazoefficiens* strain USDA110(WT) or USDA110(GUS), grown at 30°C in HM medium (Cole and Elkan, 1973) supplemented with appropriate antibiotics for 3 days. After incubation, bacterial cultures were centrifuged at 5,000 × g for 10 min, washed twice with sterile DI water, and resuspended to an OD600 of 0.1 in sterile DI water for inoculation. Plants were grown in nitrogen-free B&D medium (Broughton and Dilworth, 1971) or supplemented with 0.5 mM NH_4_NO_3_ for non-infected controls. Root samples were harvested at designated time points in a sequential manner, ensuring that nodules were carefully removed prior to collection. The harvested roots were immediately frozen in liquid nitrogen and stored at −80°C for further analysis. All experiments included three biological replicates.

### Cloning and Plasmid Construction

Promoter sequences of *Glyma.01G040700* and target genes were amplified from Williams 82 genomic DNA. The promoter, *GAL4-VP16*, nos terminator and *5xUAS* were cloned into *pSoyGUS* (Hossain et al., 2020)(KpnI, PstI digested) using Gibson assembly. Likewise, the promoter fragment, *GAL4-VP16*, nos terminator, *5xUAS*, and *HF (His-Flag)-GmRPL18* were assembled into *pCAMGFP-CvMV-GWi binary vector* (Graham et al., 2007) (KpnI, PstI digested) using Gibson assembly. All primers used for cloning are listed in Supplementary Table 8.

For Ox constructs, *pCAM-Ruby-Ox* plasmid was generated by linearizing *pSoyGUS* with PstI and SacII, followed by Gibson Assembly with PCR-amplified fragments of the Ruby gene from the *35S:RUBY* plasmid (He et al., 2020), and the *Ox* promoter from a corresponding template. The coding sequence of *GmbHLH121* (*Glyma.03G105700*) was amplified from cDNA templates and cloned into PstI site by Gibson assembly.

For RNA interference (RNAi) constructs, *pCAMRUBY-CvMV-GWi* binary vector was newly established by cloning the gateway RNA interference (GWi) sequences into PstI site of *pCAM-Ruby-Ox* vector. Approximately 160bp gene-specific fragment of *GmbHLH121* was amplified using cDNA and cloned into *pDONR/Zeo*, followed by recombination into the *pCAMRUBY-CvMV-GWi* binary vector. For the Promoter::GUS construct, approximately 2.0 kb sequences of *GmbHLH121* promotor were amplified from soybean gDNA and cloned by Gibson assembly into the *pSoyGUS* vector (KpnI, PacI digested).

### Soybean Hairy Root Transformation and Microscopy

*Agrobacterium rhizogenes* strain K599 carrying target constructs was used for hairy root transformation as described previously(Song et al., 2021). Transgenic roots were selected based on GFP fluorescence using a dissecting fluorescence microscope or Ruby color by visual inspection before and after inoculation with *B. diazoefficiens*. Nodule numbers were recorded 4 weeks post-inoculation. Harvested roots were flash-frozen in liquid nitrogen for molecular analysis. For histological studies, nodules were hand-sectioned and subjected to GUS staining before visualization. Stained sections were examined using bright-field microscopy (Leica DM5500B). Statistical analysis included at least 10 plants per replicate, and Student’s t-test or ANOVA followed by Tukey’s HSD test were used for significance testing.

### Translating Ribosome Affinity Purification (TRAP) and RNA Sequencing (TRAP-Seq)

TRAP was performed using His-FLAG-tagged GmRPL18 as described by Zanetti et al. (2005) with modifications (Song et al., 2022). Phloem-enriched hairy root tissues expressing HF-GmRPL18 were harvested at 72 hpi and 21 dpi. Approximately 10 g of root material per sample was used for immunoprecipitation, yielding at least 400 ng of mRNA for library construction. Three biological replicates were analyzed per time point.

### RNA-Seq Library Preparation and Sequencing

Total RNA extracted from TRAP samples was shipped on dry ice to the University of Missouri DNA Core Facility. Libraries were prepared using the TruSeq Stranded mRNA Kit (Illumina) and sequenced (75 bp paired-end) on an Illumina NextSeq 500 platform.

### Bioinformatics Analysis of TRAP-Seq Data

FASTQ files were quality-checked using MultiQC (version 1.9) (Ewels et al., 2016), and low-quality reads were filtered. High-quality reads were mapped to the *Glycine max Wm82.a4.v1* genome using STAR (version 2.7.4a) (Dobin et al., 2013), and a raw expression matrix was generated using HTSeq (version 0.14) (Anders et al., 2015). Differential expression analysis was performed with edgeR (version 3.32.1) (Robinson et al., 2010). Heatmaps were generated using log2 fold-change values for significant genes. The identified DEGs were analyzed and annotated using PlantTFDB v5.0 (Tian et al., 2020) to identify TFs within the DEGs. RNA-seq data were deposited into the NCBI GEO database (GSE292049).

### Functional and Pathway Enrichment Analysis

Gene Ontology (GO) enrichment analysis was performed using the tool described by (Morales et al., 2013) and results were further summarized using Revigo(Supek et al., 2011) to remove redundant GO terms. KEGG pathway analysis was conducted using g:Profiler (Raudvere et al., 2019). All analyses were conducted using default parameters unless otherwise specified.

### Cis-Regulatory Element Analysis

Cis-element analysis was conducted on 2 kb promoter sequences using PlantCARE (Lescot et al., 2002). Promoter sequences were submitted to the PlantCARE database to identify regulatory motifs. The analysis included scanning for hormone-responsive elements, stress-related regulatory sites, and transcription factor binding motifs. All parameters were set to default unless otherwise specified.

### Histochemical GUS Staining

GUS staining was conducted following (Jefferson et al., 1987). Transgenic roots were incubated overnight at 37°C in GUS staining solution (1 mg/mL X-Gluc, 100 mM sodium phosphate buffer, pH 7.0, 1 mM EDTA, 0.05% Triton X-100, 5 mM potassium ferrocyanide, 5 mM potassium ferricyanide). Stained roots were imaged using a Leica DM5500B microscope.

### Quantitative Real-Time RT-PCR (qRT-PCR)

Total RNA was extracted from transgenic roots and treated with DNase I. cDNA was synthesized from 2 μg of total RNA using M-MLV Reverse Transcriptase (Promega). qRT-PCR was performed using SYBR Green PCR mix (ABI) on a Bio-Rad CFX96 system. Relative transcript levels were normalized against the Cons6 reference gene (Libault et al., 2008). Primer sequences are listed in Supplementary Table 10.

### Immunoprecipitation and Western Blot Analysis

Total proteins were extracted from approximately 200 mg of transgenic root tissues. For immunoprecipitation, 1 mg of protein extract was incubated with ANTI-FLAG M2 Affinity Gel (Sigma) following the manufacturer’s protocol. Western blotting was performed using monoclonal anti-FLAG-HRP antibody (1:3000, Sigma). Detection was carried out using enhanced chemiluminescence (ECL).

## Author Contributions

J.S. and G.S. conceived and designed the study. J.S. and L.S. conducted the bioinformatics analysis, while J.S. and B.L. were responsible for sample collection. J.S., S.A., S.T., and Y.C. carried out the experiments. D.X. revised the manuscript. J.S. and S.A. wrote the manuscript with input from G.S. All authors read and approved the final version.

## Notes

### Competing Interest Statement

The authors have declared no competing interest.

