## Supplementary Figures for "Phloem-Specific Translational Regulation of Soybean Nodulation: Insights from a Phloem-Targeted TRAP-Seq Approach"

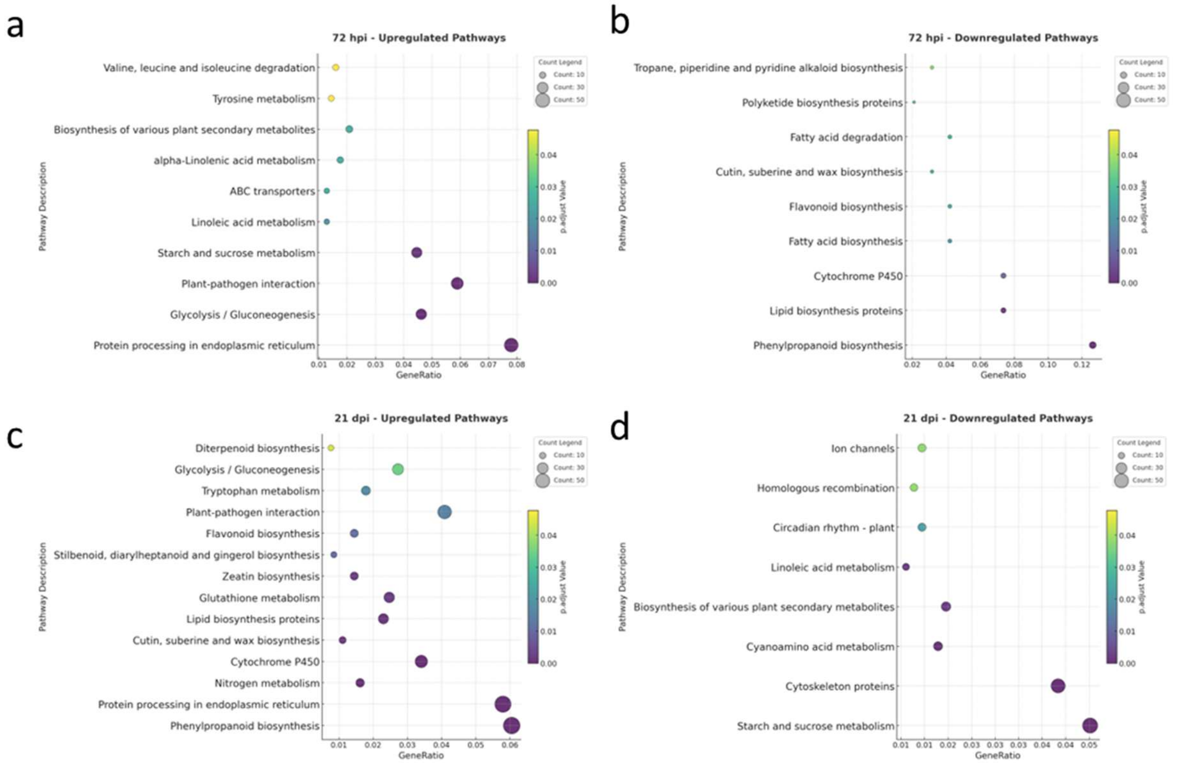


**Figure S1. KEGG Pathway Enrichment Analysis of Differentially Expressed Genes in Soybean Root Phloem**

KEGG pathway enrichment analysis of DEGs in soybean root phloem at 72 hpi and 21 dpi. Each bubble represents an enriched KEGG pathway, with bubble size corresponding to the number of DEGs mapped to the respective pathway and color intensity denoting statistical significance (-log10(P-value)). The x-axis represents the GeneRatio, defined as the proportion of DEGs associated with a given pathway relative to the total DEGs analyzed. Pathways are categorized based on expression patterns: (a) upregulated pathways at 72 hpi, (b) downregulated pathways at 72 hpi, (c) upregulated pathways at 21 dpi, and (d) downregulated pathways at 21 dpi. This analysis provides insights into dynamic transcriptional reprogramming within the root phloem in response to rhizobacterial symbiosis.

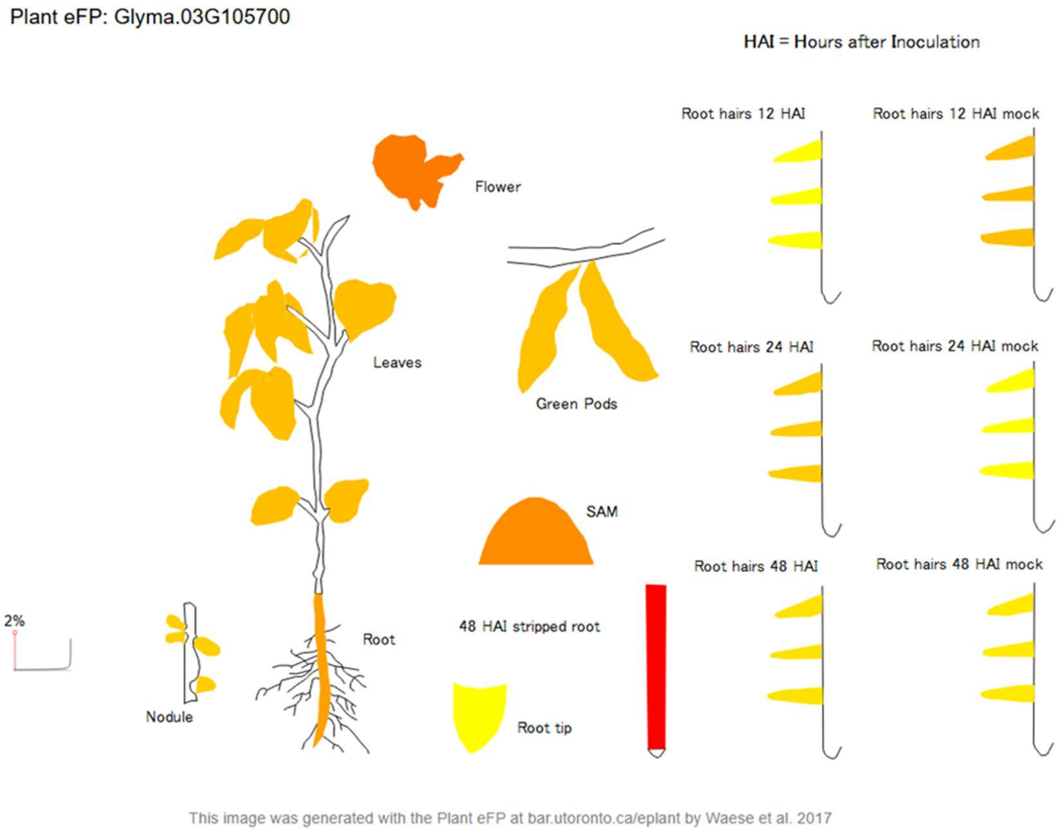


**Figure S2. Spatial Expression Profile of *Glyma.03G105700* in Various Soybean Tissues**

Expression distribution of *Glyma.03G105700* across various soybean tissues and developmental stages, visualized using the Plant eFP Browser. Gene expression levels are represented by a color gradient from yellow (low expression) to red (high expression), with grayscale indicating masked data (≥100% RSE, relative standard error).

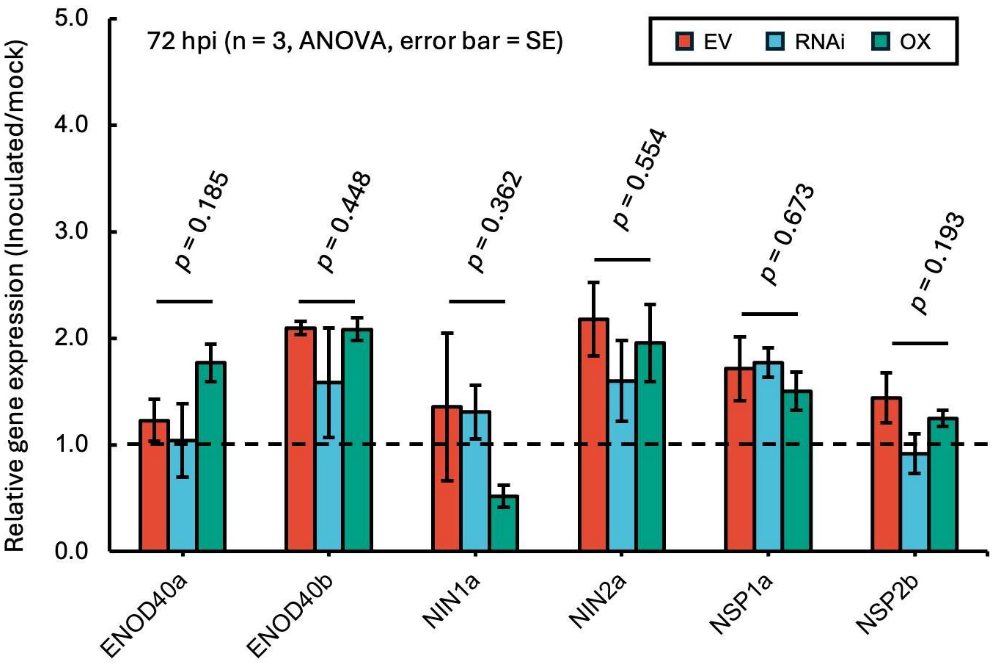


**Figure S3. qRT-PCR Analysis of Early Nodulation-Related Genes in *GmbHLH121* RNAi and Overexpression (OX) Hairy Roots**

Relative expression levels of early nodulation-related genes were assessed in empty vector (EV), *GmbHLH121* RNA interference (RNAi), and *GmbHLH121* overexpression (OX) hairy roots using quantitative reverse transcription PCR (qRT-PCR). Gene expression was normalized against the reference gene *Cons6*, and fold changes were calculated relative to the EV control. Error bars represent standard error.
